# RAB6 GTPase is a crucial regulator of the mammary secretory function controlling STAT5 activation

**DOI:** 10.1101/2020.03.12.989236

**Authors:** Surya Cayre, Marisa M. Faraldo, Sabine Bardin, Stéphanie Miserey-Lenkei, Marie-Ange Deugnier, Bruno Goud

## Abstract

The Golgi-associated RAB GTPases, RAB6A and RAB6A’, regulate anterograde and retrograde transport pathways from and to the Golgi. *In vitro*, RAB6A/A’ have been reported to be involved in several cellular functions, including, in addition to transport, cell division, migration, adhesion and polarity. However, their role remains poorly described *in vivo*, in particular in epithelial tissues. Here, we generated BlgCre; *Rab6a*^F/F^ mouse presenting a specific deletion of *Rab6a* in the mammary luminal secretory lineage during gestation and lactation. *Rab6a* loss severely impaired the differentiation, maturation and maintenance of the secretory tissue, compromising lactation. It led to a decreased activation of STAT5, a key regulator of the lactogenic process primarily governed by prolactin. Data obtained with a human mammary epithelial cell line suggested that defective STAT5 activation might originate from a perturbed transport of the prolactin receptor, altering its membrane expression and signaling cascade. Despite the major functional defects observed upon *Rab6a* deletion, the polarized organization of the mammary epithelial bilayer was preserved. Altogether, our data reveal a crucial role for RAB6A/A’ in the lactogenic function of the mammary gland. They also suggest that the trafficking pathways controlled by RAB6A/A’ depend on cell type specialization and tissue context.

**SUMMARY STATEMENT:** This study reveals a role for the Golgi-associated RAB GTPases, RAB6A/A’, in the lactogenic function of the mammary gland.

## INTRODUCTION

Small GTPases of the RAB family are master regulators of vesicular transport and membrane trafficking in eukaryotic cells. Through their specific intracellular localization and ability to recruit various types of effectors, RAB GTPases tightly control the molecular exchanges between cell compartments (Stenmark, 2009). This large family comprises about 70 members in human and includes four RAB6 proteins, RAB6A and its splice variant RAB6A’, RAB6B and RAB6C. The ubiquitously expressed RAB6A/A’ are the most abundant Golgi-associated RAB GTPases and have been reported to control anterograde and retrograde trafficking from and to the Golgi apparatus (Goud *et al.*, 2018).

RAB6A/A’ has been intensively studied in cultured cells and implicated in several cellular functions, including cell division, migration, adhesion, polarity establishment and secretory mechanisms (Miserey-Lenkei *et al.*, 2006 and 2010; Micaroni *et al.*, 2013; Shafaq-Zadah *et al.*, 2016; Fourrière *et al.*, 2019; Homma *et al.*, 2019). Importantly, we recently reported that RAB6A/A’ are essential for early embryonic development. RAB6A/A’ null mouse embryos die at day 5.5, exhibiting defective adhesion between the epiblast layer and the visceral endoderm, with a disorganization of the basement membrane and a perturbed expression of the β1 integrin chain in epiblast cells (Shafaq-Zadah *et al.*, 2016).

To investigate the role of RAB6A/A’ in specific tissular context, we have generated *Rab6a*^F/F^ mice carrying a conditional null allele of *Rab6a* that allows the targeted deletion of RAB6A/A’ (Bardin *et al.*, 2015). Using this model, we have shown that RAB6A/A’ is required for the proper maturation of melanosomes in melanocytes and that they play a key role in T lymphocyte activation (Patwardhan *et al.*, 2017; Carpier *et al.*, 2018).

RAB6 function in highly polarized epithelial tissues remains poorly documented yet. Mammary gland provides a unique model to study morphogenesis, epithelial tissue polarity, cell fate specification and secretory mechanisms. Its development, mainly postnatal, comprises two distinct morphogenetic events: the growth and branching of epithelial ducts during puberty and the lobulo-alveolar development during gestation (Macias and Hinck, 2012; Brisken and Atacca, 2015). At each pregnancy, under progesterone and prolactin (PRL) stimulation, the mammary epithelium undergoes proliferation and differentiation preparing it for milk secretion, its primary function. At parturition, a fall in progesterone accompanied by elevated PRL levels triggers secretory activation and lactation (Anderson *et al.*, 2007). PRL and its receptor (PRLR) play a pivotal role in alveologenesis and lactation, in particular through activation of the transcription factor STAT5 (Hennighausen and Robinson, 2008).

In ducts and alveoli, the mammary epithelium comprises an outer layer of basal myoepithelial cells and an inner layer of luminal cells organized around a lumen. This bilayered structure sits on a basement membrane surrounded by stromal elements. During lactation, the secretory luminal cells produce milk components, including milk proteins, lactose and lipids, whereas the contractile myoepithelial cells serve for milk expulsion (Anderson *et al.*, 2007; Moumen *et al.*, 2011). Luminal cells are highly polarized, with an apical domain facing the lumen and a basolateral domain contacting adjacent luminal and myoepithelial cells. In addition, during lactation, secretory luminal cells interact with the basement membrane, myoepithelial cells forming a discontinuous layer around alveoli. Due to their specialized function and numerous interactions, luminal cells display a large set of apical and baso-lateral surface proteins (Glukhova and Streuli, 2013; Chatterjee and McCaffrey, 2014).

The mammary luminal cell population is heterogeneous, comprising hormone-sensing cells positive for estrogen and progesterone receptors (ER, PR) and cells devoid of ER/PR expression (Brisken and Ataca, 2015). The secretory lineage is largely composed of ER/PR^-^ cells. It contains distinct stem/progenitor cells that are amplified during gestation and give rise to fully functional secretory cells in the lactating gland (Rodilla *et al.*, 2015; Bach *et al.*, 2017). To study the role of RAB6A/A’ in mammary gland development and function, we generated BlgCre; *Rab6a*^F/F^ mice, in which *Rab6a* was deleted specifically in the secretory lineage via the beta-lactoglobulin (*Blg*) promoter, primarily active throughout pregnancy and lactation (Selbert *et al.*, 1998; Chapman *et al.*, 1999; Naylor *et al.*, 2005; Molyneux *et al.*, 2010; Romagnoli *et al.*, 2020).

## RESULTS

### RAB6A/A’ expression is upregulated in the luminal secretory lineage during gestation

To get insight into RAB GTPase expression in mammary luminal cells, we first analyzed their transcriptomic profiles. In the adult virgin gland, the ER/PR^-^ luminal cell population, targeted by the *Blg* promoter, can be separated from the ER/PR^+^ cell fraction using ICAM-1 expression, as described in a previous work (Di-Cicco *et al.*, 2015). In flow cytometry, ICAM-1, combined with the epithelial-specific marker CD24, discriminates CD24^low^ ICAM1^+^ basal cells, CD24^high^ ICAM1^-^ luminal cells (ER/PR^+^, referred to as HR^+^) and CD24^high^ ICAM1^+^ luminal cells (mostly ER/PR^-^; referred to as HR^-^) (Fig.1A; Di-Cicco *et al.*, 2015). Using the transcriptomic profiles of the HR^+^ (ICAM1^-^) and HR- (ICAM1^+^) luminal cell subsets that we previously established (Chiche *et al.*, 2019), we analyzed the expression levels of the 58 *Rab* GTPase genes present in the microarrays. Only 19 *Rab* genes were found robustly expressed in both HR^+^ and HR^-^ luminal cell populations (Fig. 1B). These include, in addition to *Rab6a, Rab1a/b, Rab2a, Rab4a, Rab5a/c, Rab7a, Rab8a/b, Rab9a, Rab10, Rab11a/b, Rab14, Rab18, Rab21, Rab24* and *Rab25.* HR^+^ and HR^-^ luminal cell populations displayed very similar *Rab* profiles, *Rab1a, Rab2a, Rab7a, Rab10* and *Rab14* being the top ranked genes. Of note, they did not significantly express *Rab6b* (Fig. 1B), known to be mainly expressed in neuronal and neuro-endocrine cells (Opdam *et al.*, 2000; Goud *et al.*, 2018).

**Figure 1.**
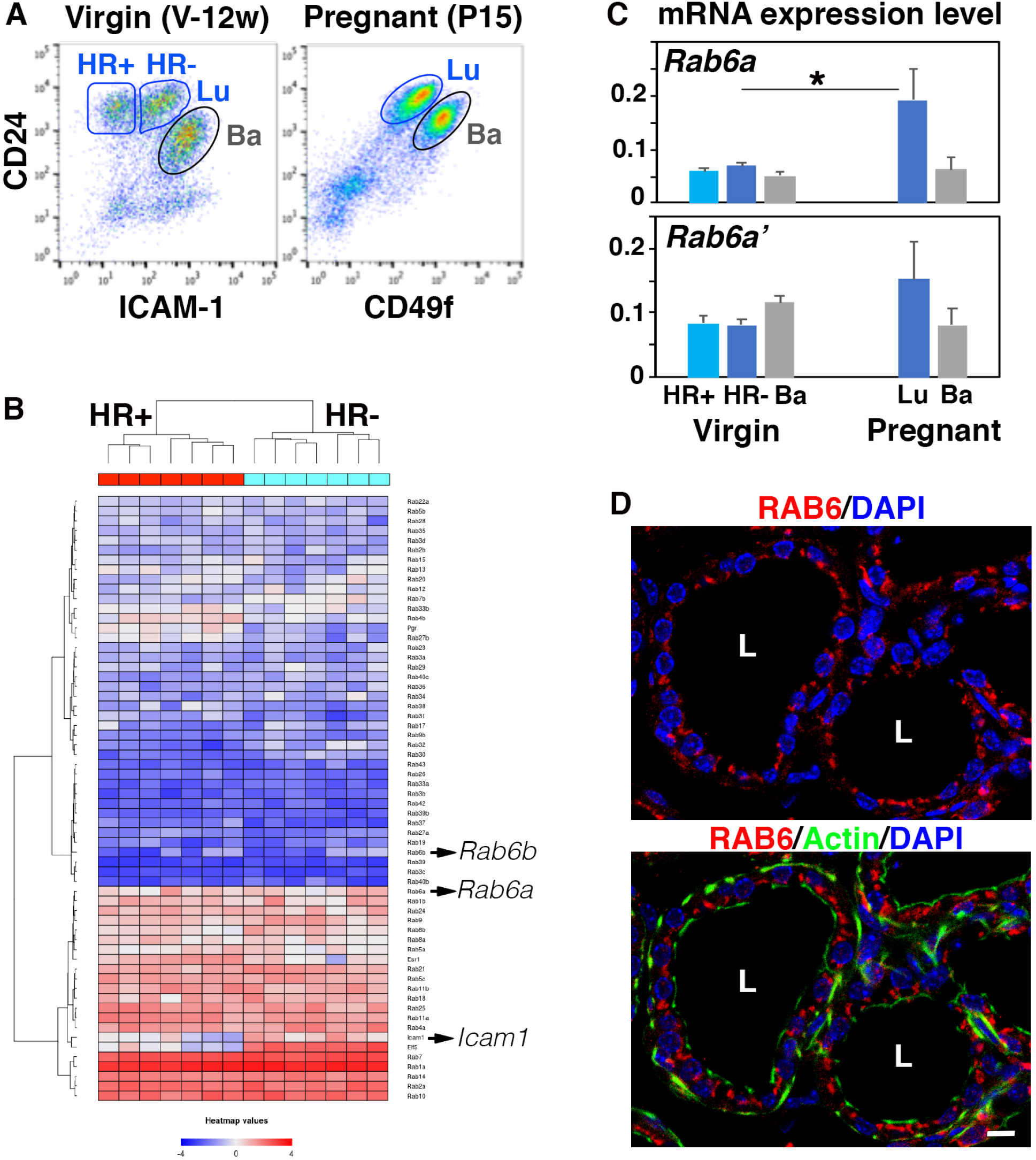
*Rab6* expression in mammary luminal and basal cells isolated from adult virgin and pregnant females. **(A)** Flow cytometry analysis of mammary cells isolated from 12-week-old virgin mice (left; V-12w) and 15-day pregnant mice (right, P15). The gated subsets within the CD24^+^ epithelial cell pool are: CD24^low^ ICAM1^+^ basal cells (Ba), CD24^high^ ICAM1^-^ and CD24^high^ ICAM1^+^ luminal cells (HR^+^ and HR^-^, respectively) (left); CD24^low^ CD49f^high^ basal cells (Ba) and CD24^high^ CD49f^low^ (right). **(B)** Heat map based on microarray data (Chiche *et al.*, 2019) showing the relative gene expression of 58 RAB GTPases in the HR^+^ and HR^-^ luminal cell populations isolated from adult virgin mice as shown in A (left). Genes expressed at a significant level appear in pink to red. The heatmap includes the reference genes, *Icam1, Elf5, Pgr* (encoding PR) and *Esr1* (encoding ER) used to characterize the luminal cell populations. Microarray data are from 7 separate sorting experiments. **(C)** qPCR analysis of *Rab6a* and *Rab6a*’ expression in mammary basal and luminal cells isolated from adult virgin and pregnant mice as shown in (A). Data are the mean ± SEM of 3-4 separate cell preparations. *p=0.039 **(D)** RAB6 immunolocalization in a lactating mammary gland. Actin marks basal myoepithelial cells and the apical pole of luminal cells facing the alveolar lumen (L). Nuclei are stained with DAPI. Bar, 10 μm.

Consistent with the microarray analysis, qPCR data confirmed that similar amounts of *Rab6a* transcripts were detected in HR^-^ and in HR^+^ luminal cells isolated from virgin glands (Fig.1C). *Rab6a*’ was also present in both luminal subsets, at level close to that of *Rab6a* (Fig.1C). The two variants were expressed in basal cells at similar level than in luminal cells (Fig.1C).

To further investigate *Rab6a* expression in the mammary epithelium, we isolated luminal and basal cells from mice at day 15 of gestation (Fig.1A), using differential expression of CD24 and α6 integrin chain (CD49f) as described (Visvader and Stingl, 2014). At this stage, the luminal population is largely composed of HR^-^ cells that have been amplified upon hormonal stimulation and are committed to the secretory lineage (Macias and Hinck, 2012). Interestingly, qPCR data showed that luminal cells from pregnant mice displayed higher levels of *Rab6a* than those from adult virgin mice (Fig.1C). *Rab6a*’ expression in luminal cells exhibited the same tendency than that of *Rab6a* (Fig.1C). *Rab6a* and *Rab6a*’ levels were not significantly modulated in basal cells upon gestation (Fig.1C).

Overall, these data demonstrate that *Rab6a/a*’ expression is upregulated during pregnancy in luminal cells only and therefore suggest an important role for RAB6A/A’ in the secretory lineage. Consistently, using an antibody recognizing all RAB6 isoforms, we observed a strong RAB6 expression in the luminal secretory cells of the lactating gland (Fig.1D). As expected for a Golgi-associated protein, RAB6 was concentrated at the apical pole of the luminal cells.

### Mammary development and epithelium organization of BlgCre; *Rab6a*^F/F^ mice are not critically affected at day 15 of pregnancy (P15)

The *Blg* promoter activity has been reported to substantially increase from midpregnancy (around P10) and culminate during lactation (Selbert *et al.*, 1998). To evaluate the role of RAB6A/A’ (hereafter referred to as RAB6A) in the development of the luminal secretory lineage, we first analyzed the mammary phenotype of BlgCre; *Rab6a*^F/F^ mutant mice at P15, using *Rab6a*^F/F^ littermates as controls. Quantitative flow cytometry analyses showed that mutant and control mammary tissues displayed similar percentages of CD24+ epithelial cells with identical luminal/basal cell ratios (Figs. S1A-B). Surface expression of α6 or β1 integrin chains was not perturbed in mutant luminal and basal cells (Fig. S1A). The extent of *Rab6a* deletion, checked in isolated mutant luminal cells by qPCR, was estimated at 80% (Fig. S1C), in agreement with previous studies using BlgCre-mediated gene deletion (Chapman *et al.*, 1999; Naylor *et al.*, 2005; Romagnoli *et al.*, 2020).

In line with the flow cytometry data, whole-mount analyses did not reveal differences between mutant and control mammary trees that displayed similar fat pad occupancy and branching patterns (Fig. S1D). Organization of the mammary bilayer, evidenced by double labeling for the luminal- and basal-specific keratins, K8 and K5, was not perturbed in mutant mice (Fig. S1D). As expected at this developmental stage, mutant and control mammary epithelium were actively proliferating (Fig. S1D). They both displayed typical alveolar buds with nascent cytoplasmic lipid droplets (CLD) positive for adipophilin (ADPH) (Fig. S1E), a major CLD protein whose expression is specifically induced in alveolar luminal cells around mid-pregnancy (Anderson *et al.*, 2007; Russel *et al.*, 2011). Percentages of PR^+^ luminal cells were similar in mutant and control epithelium (Fig. S1F), indicating that the relative proportion of HR^-^ and HR^+^ luminal lineages was not affected upon *Rab6a* deletion.

Collectively, these data indicated that *Rab6a* deletion in luminal cells did not critically impact the first set of pregnancy-associated developmental events, resulting in amplification of basal and luminal cell populations, side branching and alveolar bud formation. Moreover, the bilayered organization of the mammary epithelium and expression of luminal cell markers, such as K8, ADPH, α6 and β1 integrin chains, were preserved upon *Rab6a* deletion.

### BlgCre; *Rab6a^F/F^* mice exhibit altered alveolar differentiation at late gestation

We next analyzed the mammary phenotype of BlgCre; *Rab6a*^F/F^ mutant mice at day 18 of pregnancy (P18). This prelactational stage is characterized by well described histological changes and secretory differentiation events. These include alveolar distension with luminal space enlargement, presence of large CLDs in alveolar luminal cells and substantial milk protein expression, in particular caseins, Whey Acidic Protein (WAP) and a-lactalbumin, an essential cofactor for milk lactose synthesis (Oakes *et al.*, 2006; Anderson *et al.*, 2007; Oakes *et al.*, 2017).

Whole-mount images indicated that the mutant epithelium, although well developed, appeared less dense than the control (Fig. 2A). In line with this observation, histological analyses of hematoxylin/eosin-stained sections showed that the alveoli from mutant glands were condensed compared to those from control glands (Fig. 2B), evoking reduced amounts of secretory products. Quantification of alveolar size distribution confirmed that unlike controls, the mutant glands contained a majority of alveoli of small to medium size and almost no large alveoli (Fig. 2C). This phenotype was not associated with reduced proliferation capacities. Indeed, proliferation index was twice as high in the mutant compared to the control epithelium (Fig. 2D), indicating that RAB6A-deficient mammary glands, unlike controls, were still in an active growing phase.

**Figure 2.**
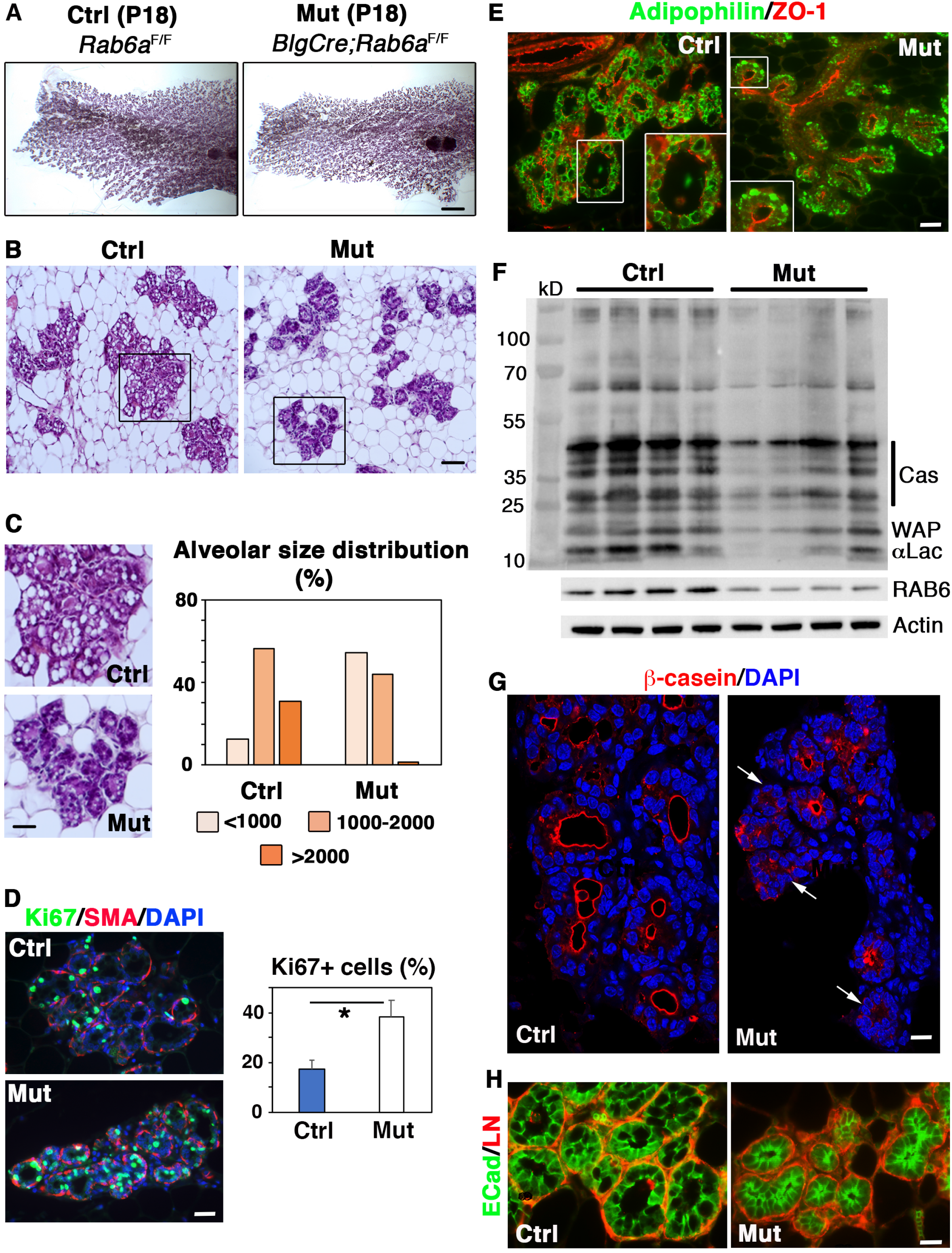
Impaired alveolar luminal cell differentiation in Blg-Cre; *Rab6a*^F/F^ females at late pregnancy (P18) **(A)** Carmine-stained whole-mounts of mammary glands from control and mutant females at P18. Bar, 2.25 mm. **(B)** Views of hematoxylin/eosin-stained histological sections through control and mutant mammary glands at P18. Bar, 100μm. **(C)** Left: Enlarged views of control and mutant alveoli shown in (B) Bar, 48μm. Right: compared distributions of alveolar size at P18, estimated in pixel. Data are from a pool of measurements performed on two distinct histological sections from 3 control and 3 mutant mammary glands. A total of 1450 alveoli was measured. Pearson’s Chi-square test: p<0.001 **(D)** Cell proliferation in control and mutant mammary glands at P18, estimated by Ki67 expression. Left: triple Ki67/SMA/DAPI immunofluorescent staining. SMA (smooth muscle actin) marks basal myoepithelial cells surrounding alveoli. Bar, 35μm. Right: Percentages of Ki67+ luminal cells. Data are the mean ± SEM of counts performed on sections though 3 distinct control and mutant mammary glands. About 2000 DAPI-stained nuclei were counted on each section. *p<0.001 **(E)** Immunolocalization of adipophilin and ZO-1 in control and mutant mammary tissues at P18, with enlarged views of individual alveoli. Bar, 45μm. **(F)** Western blots for milk proteins, RAB6 and actin (used as loading control) performed with 4 distinct control and mutant mammary gland protein extracts. Molecular size markers are shown on the left and the corresponding milk proteins, α-lactalbumin (aLac), whey acidic protein (WAP) and caseins (Cas) on the right. Quantification of RAB6 depletion is shown in Fig. S2A. **(G)** Immunolocalization of β-casein in control and mutant mammary epithelium at P18. Arrows indicate the mutant alveoli devoid of b-casein expression. Bar, 20μm. **(H)** Double immunolabeling of E-cadherin and laminin (ECad/LN; bar, 15μm) in control and mutant mammary tissues at P18.

Notably, hematoxylin/eosin staining revealed that alveoli from the mutant tissue displayed a poor number of CLDs (Fig. 2C). Immunofluorescent studies confirmed the presence of large ADPH-coated CLDs in alveolar luminal cells from control glands (Fig. 2E). In contrast, mutant alveolar luminal cells primarily contained small ADPH-expressing CLDs often located at the basal pole (Fig. 2E), a distribution pattern observed at P15 in both control and mutant tissues (Fig. S1E). Thus, RAB6A deficiency appeared to impair or considerably delay CLD maturation, indicating perturbed secretory differentiation events.

To further analyze the differentiation features characterizing late gestation, we performed western blotting (WB) on mammary gland extracts using an antibody against mouse milk-specific proteins, as reported (Oakes *et al.*, 2017). The data showed that the mutant samples contained lower amounts of milk components than the controls, including α-lactalbumin, caseins and WAP (Fig. 2F). Efficiency of RAB6 depletion was controlled in parallel using an antibody recognizing RAB6A/A’ and RAB6B (Fig. 2F and S2A). In line with the WB data, immunolocalization studies showed that in the mutant tissue, numerous alveoli lacked β-casein expression whereas in controls, almost all alveoli displayed a strong β-casein staining lining the lumen (Fig. 2G).

Immunodetection of lineage-specific markers showed that overall, the polarized organization of the mutant mammary epithelium was preserved at late pregnancy. In mutant as in control alveoli, K8-positive luminal cells were surrounded by basally-located myoepithelial cells expressing K5 and smooth-muscle actin (SMA) (Fig. S2B). Laminin, a major component of the basement membrane, appeared normally deposited around the alveoli of the mutant tissue (Fig. 2H). Major markers of apical and baso-lateral polarity, such as ZO-1, MUC1, E-cadherin and β1 integrin, were strongly expressed and properly localized in the mutant alveolar luminal cells (Fig. 2H and S2B-D). ZO-1 and MUC1 staining well illustrated the reduced luminal space of the condensed mutant alveoli (Fig. S2C). Flow cytometry analyses confirmed that surface expression of α6 and β1 integrin chains was not perturbed in the mutant epithelium, neither in luminal nor in basal cells (Fig. S2E).

Together, the data obtained at late pregnancy revealed impaired or delayed differentiation events in BlgCre; *Rab6a*^F/F^ mammary glands. In particular, the mutant epithelium exhibited deficient CLD maturation and decreased milk protein production, most probably accounting for the defective alveolar enlargement. This indicated an important role for RAB6A in the alveolar differentiation stage characterizing the prelactational phase. Noticeably, perturbations induced by loss of *Rab6a* occurred without any obvious alterations of the bilayered organization of the mammary epithelium and the apico-basal polarity of the alveolar luminal cells, suggesting functional rather structural defects.

### BlgCre; *Rab6a*^F/F^ mice display a severe lactation defect

In primiparous control mice, at day one of lactation (L1), the lobulo-alveolar structures almost entirely occupied the mammary fat pad and the milk-producing alveoli were markedly dilated, as seen on whole-mounts and histological sections (Fig. 3A). In contrast, the mammary epithelium from BlgCre; *Rab6a*^F/F^ mutant mice showed a significant reduction in fat pad occupancy (Fig. 3A) and as observed at late gestation, the mutant tissue was devoid of large alveoli (Fig. 3A and S3A). Immunohistofluorescence studies showed that as expected, RAB6 was absent from the secretory luminal cells of BlgCre; *Rab6a*^F/F^ mice at L1 whereas it was detected in the non-targeted basal myoepithelial cells (Fig. 3B). Alveolar luminal cells lacking RAB6 exhibited a normal distribution of GM130, a cis-Golgi marker preferentially localized at their apical pole (Fig. 3B). The bilayered organization of the mutant mammary epithelium was conserved and although poorly distended, alveoli were decorated with myoepithelial cells displaying a stellate shape, typical of the lactation period (Fig. 3C). Mutant as control alveoli were enveloped by a continuous basement membrane rich in laminin (Fig. 3D).

**Figure 3.**
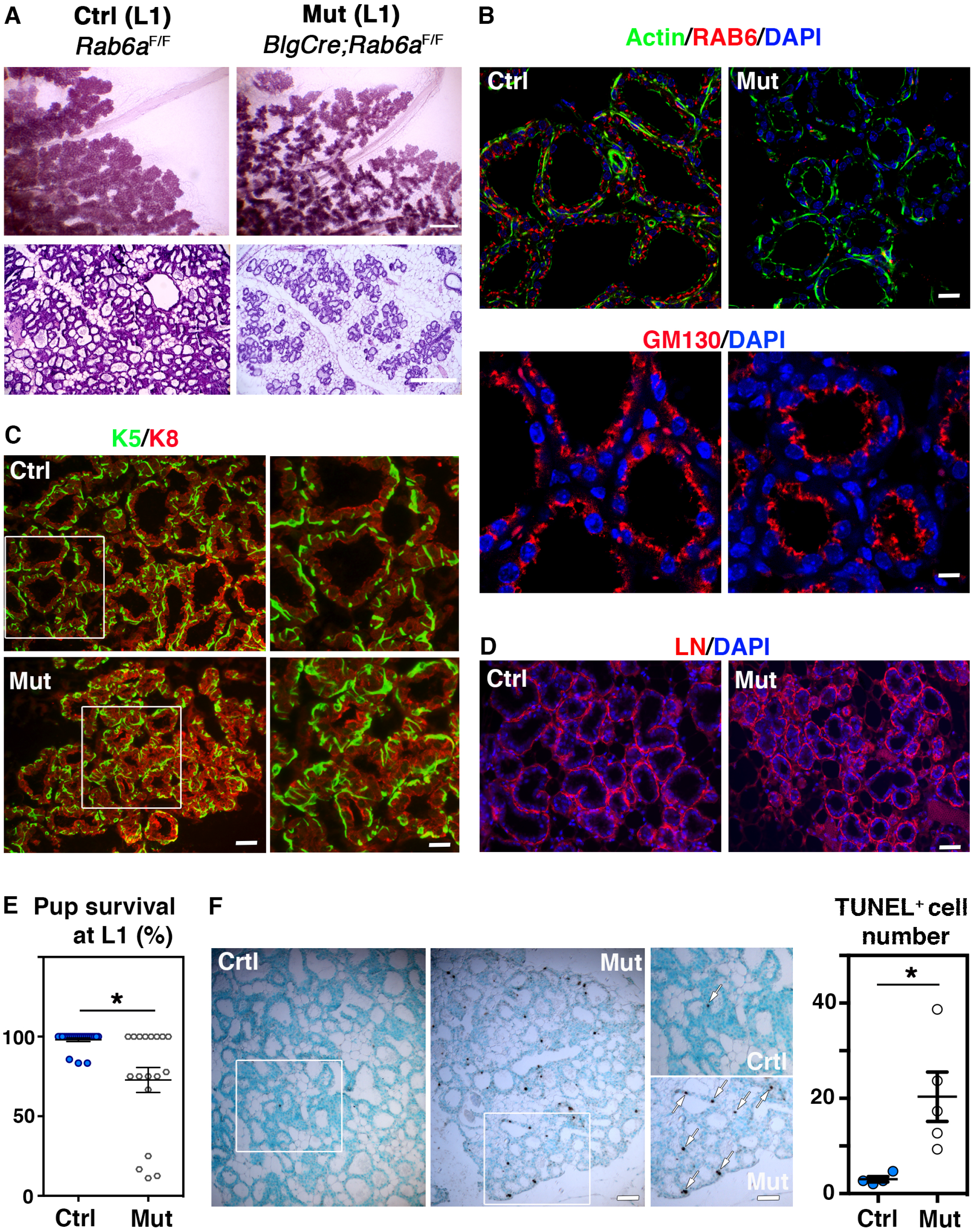
Mammary phenotype and lactation deficiency of Blg-Cre; *Rab6a*^F/F^ females at day 1 of lactation (L1) **(A)** Carmine-stained whole-mounts (upper panels; bar, 0.3 mm) and hematoxylin/eosin-stained histological sections (lower panels; bar, 300μm) of mammary glands from control and mutant females at L1. **(B)** RAB6 and GM130 immunodetection in control and mutant alveolar luminal cells at L1. Upper panels: triple actin/RAB6/DAPI staining. Actin marks basal myoepithelial cells and the apical membrane of luminal cells. RAB6 is absent from mutant luminal cells but detected in basal myoepithelial cells. Bar, 20μm. Lower panels: double GM130/DAPI staining showing similar apical Golgi distribution in control and mutant luminal cells. Bar, 12μm. **(C)** Immunolabeling of basal-specific (K5) and luminal-specific (K8) keratins in control and mutant mammary epithelium at L1. Right panels: enlarged views of the delineated areas. Basal myoepithelial cells display a typical stellate shape in both control and mutant samples. Bars, 40μm (left), 20μm (right). **(D)** Immunodetection of laminin (LN) in control and mutant alveoli at L1 showing similar deposition. Bar, 75μm. **(E)** Survival of pups nursed by control and mutant primiparous dams at L1. Litters from 24 control and 18 mutant females were analyzed. *p=0.05 **(F)** Cell apoptosis in control and mutant mammary epithelium at L1, as revealed by TUNEL assay. Left panels: Views of sections through control and mutant glands. Methyl green was used as counterstaining. Enlarged views of the delineated areas are shown on the right. Arrows indicate TUNEL-positive cells. Bars, 75μm (left) and 55μm (right). Right panel: Number of TUNEL-positive cells per low-magnification microscopic field. Data are shown as mean ± SEM from four control and five mutant mice. Three distinct fields per mouse were quantified. *p=0.03

We next investigated the ability of BlgCre; *Rab6a*^F/F^ females to support growth of a first litter. Whereas almost all control dams were able to feed complete litters, eight out of the eighteen (44.5%) mutant dams analyzed had lost pups at L1 (Fig. 3E). Non-viable newborns were found to lack milk in their stomach, suggesting inability of mutant mothers to feed normally. Beyond L1, ethical reasons led us to eliminate most of the pups nursed by mutant dams, their survival being compromised by malnourishment. At L6, only four out of the eleven (36.4%) mutant dams examined were still able to feed pups (Fig. S3B). Two of them were analyzed and although nursing 3 and 5 pups respectively, they exhibited a deeply altered lobuloalveolar development (Fig. S3C). Moreover, their pups weighted significantly less than those nursed by control mothers (Fig. S3D).

As perturbed lobulo-alveolar development and lactation defects are known to trigger early cell death, we analyzed apoptosis rates in mutant and control mammary tissues at L1, using TUNEL assays. The data revealed that unlike controls, the RAB6A-depleted tissue sections contained numerous TUNEL-positive luminal cells (Fig. 3F). Apoptotic luminal cells were distributed throughout the mutant secretory tissue.

Collectively, these data showed that loss of *Rab6a* in the luminal secretory lineage severely compromised lactation and induced early cell death, in primiparous females as early as postpartum day 1.

### Secretory activation is impaired at the onset of lactation in BlgCre; *Rab6a*^F/F^ mice

Transition from pregnancy to lactation is marked by changes in the size and cellular distribution of the luminal CLDs that are considered as signs for a proper secretory activation (Anderson *et al.*, 2007). ADPH staining performed in control mammary glands at L1 showed that as expected, the large CLDs visible at late pregnancy have been replaced by numerous smaller CLDs targeted to the apical surface of the luminal secretory cells (Fig. 4A). Unlike controls, mutant luminal cells displayed CLDs of various size, including large ones that were stored in the cytosol (Fig. 4A). These data strongly suggest a failure in secretory activation in RAB6A-depleted alveolar luminal cells.

**Figure 4.**
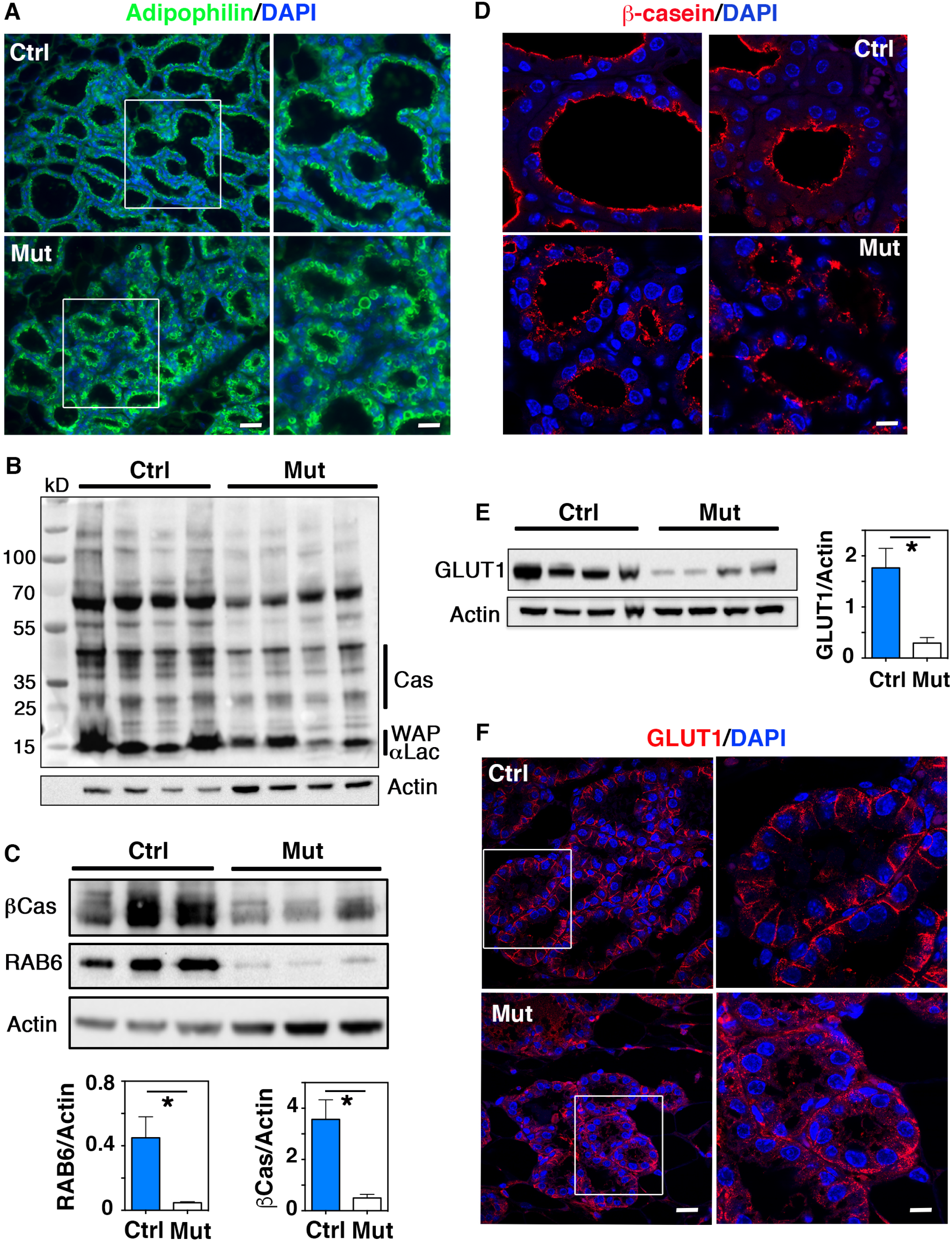
Defective post-partum secretory activation of alveolar luminal cells in Blg-Cre; *Rab6a*^F/F^ females. **(A)** Immunolocalization of adipophilin in control and mutant mammary epithelium at L1. Right panels: enlarged views of the delineated areas. Adipophilin-coated CLDs are retained in the cytosol of mutant alveolar luminal cells. Bars, 30μm (left), 15μm (right). **(B)** Western blots for milk proteins and actin (used as loading control) in 4 distinct control and mutant mammary gland extracts at L1. Molecular size markers are shown on the left and the corresponding milk proteins, αLac, WAP and caseins on the right. **(C)** Western blots for β-casein (βCas), RAB6 and actin in 3 distinct control and mutant mammary gland extracts at L1. The lower panels show the related quantifications (mean ± SEM) normalized on actin amounts. *p=0.05 (RAB6), *p=0.026 (βCas) **(D)** Immunolocalization of β-casein in control and mutant mammary epithelium at L1. Two representative fields are shown. Bar, 14μm. **(E)** Western blots for GLUT1 and actin in the 4 control and mutant mammary gland extracts analyzed in (B). The lower panel shows GLUT1 quantification (mean ± SEM) normalized on actin amounts. *p=0.026 **(F)** Immunolocalization of GLUT1 in control and mutant mammary epithelium at L1. Right panels: enlarged views of the delineated alveoli. In control, GLUT1 is sharply expressed at the baso-lateral membrane of alveolar luminal cells whereas it displays a heterogeneous and diffuse expression pattern in mutant. Bars, 20μm (left), 8μm (right).

As observed at late gestation, WB analysis at L1 showed that compared to control, the mutant mammary tissue contained lower amounts of milk proteins (Fig. 4B). Blotting with a specific anti-β-casein antibody confirmed the reduced amount of this milk protein in the RAB6A-depleted tissue samples analyzed (Fig. 4C). Immunolocalization studies showed that alveolar luminal cells from the control tissue homogeneously displayed β-casein at their apical pole. In contrast, mutant alveolar luminal cells either expressed β-casein at low level or exhibited a poorly polarized distribution pattern (Fig. 4D)

In addition to proteins and lipids, lactose is an essential milk component. Its massive synthesis from glucose during lactation is accompanied by the upregulated expression of the glucose transporter, GLUT1, at the baso-lateral membrane of alveolar luminal cells (Boxer *et al.*, 2006; Anderson *et al.*, 2007). WB analysis at L1 revealed a reduced amount of GLUT1 in the RAB6A-depleted mammary samples compared to the controls (Fig. 4E). As expected, GLUT1 was homogeneously expressed in the control tissue and sharply localized at the basolateral membrane of the alveolar luminal cells (Fig. 4F). In contrast, GLUT1 expression in mutant alveolar luminal cells was heterogeneous, with a frequent intracellular accumulation (Fig. 4F). Other polarity markers, including baso-lateral (E-cadherin, β1 integrin) and apical (ZO-1, MUC1) markers, appeared normally distributed in RAB6A-depleted luminal cells (Fig. S4A-C).

Collectively, our data revealed a postpartum failure in secretory activation of alveolar luminal cells. The reduced milk protein production, CLD retention and decreased GLUT1 expression suggest that milk amount and composition might be both affected in the absence of RAB6A, compromising pup growth and viability.

### *Rab6a* loss in alveolar luminal cells affects RAB18 expression

As described above, alveolar luminal cells from BlgCre; *Rab6a*^F/F^ females display deficient CLD maturation and apical targeting (Figs. 2E and 4A). RAB18 has been reported to control CLD growth and secretion in various lipogenic cell types (Xu *et al.*, 2018; Dejgaard and Presley, 2019). Consistently, RAB18 belongs to the RAB GTPases highly expressed in mammary luminal cells (Fig. 1B). We thus examined whether *Rab6a* loss affected RAB18 expression. Using WB, we found that RAB18 amounts were significantly decreased in the protein extracts from P18 and L1 mutant glands, compared to controls (Fig. 5A). Noticeably, the amount of other abundantly expressed RAB GTPases (RAB5, RAB8 and RAB11) was not significantly altered in the mutant samples, indicating that *Rab6a* loss had a particular impact on RAB18 (Fig. 5B).

**Figure 5.**
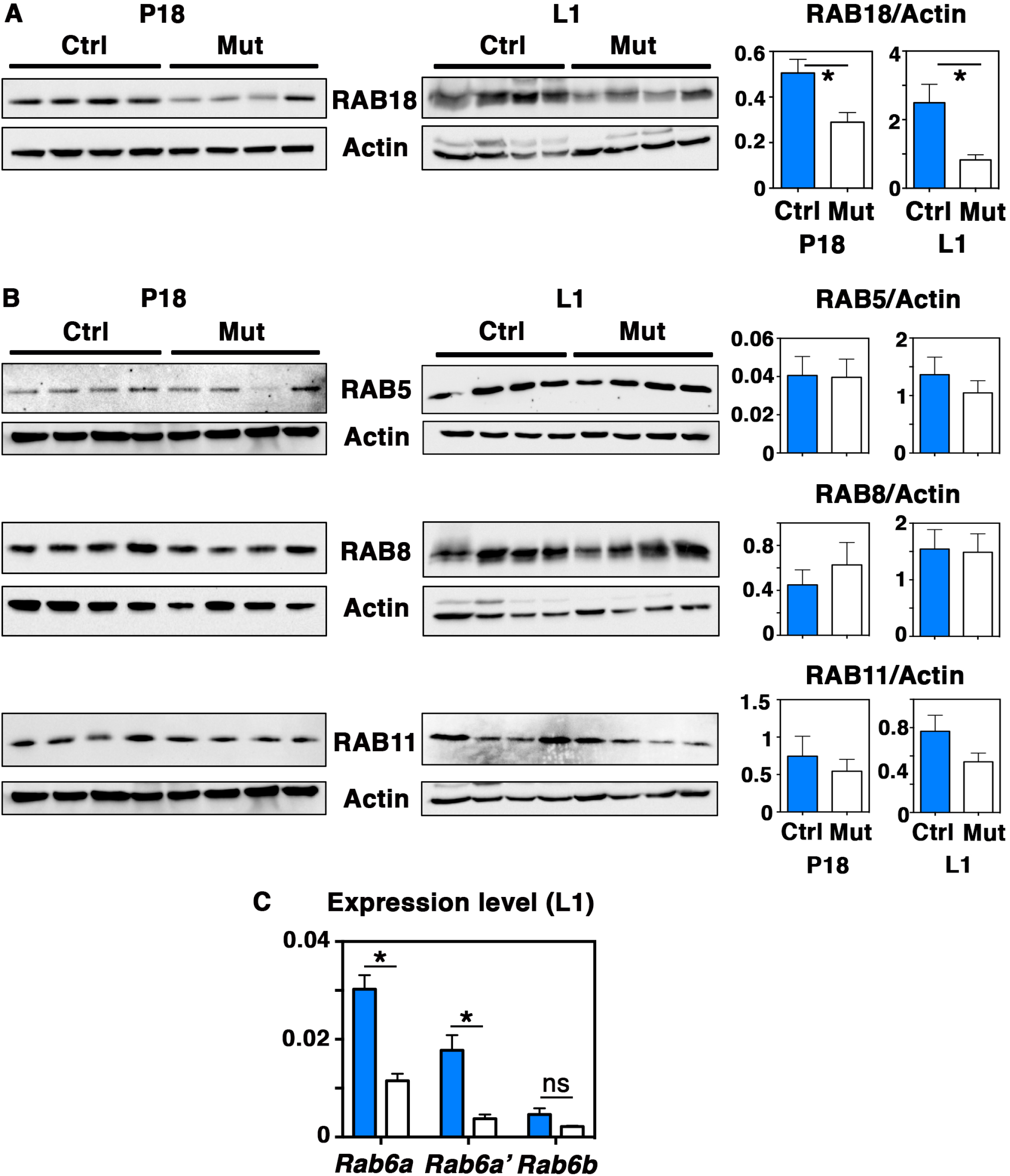
RAB expression in *Rab6a*-deficient mammary glands. **(A)** Western blots for RAB18 and actin at P18 and L1 (left panels) and quantifications (mean ± SEM) normalized on actin amounts (right panels) Data are from four distinct control and mutant mammary gland protein extracts at P18 and L1. *p=0.029 (P18), *p=0.048 (L1) **(B)** Western blots for RAB5, RAB8, RAB11 and their actin loading controls (left panels). The samples are the same than those shown in (A). The related quantifications shown on the right panels did not reveal any significant differences between control and mutant samples. **(C)** qPCR analysis of *Rab6a, Rab6a*’ and *Rab6b* mRNA expression in mammary cells derived from control and mutant glands at L1. qPCR values were normalized on *Gapdh.* Data are the mean ± SEM of 4 separate cell preparations. ns, not significant. *p=0.001 (*Rab6a*), *p=0.005 (*Rab6a’*)

In addition, using qPCR, we found that *Rab6a*-deleted and control mammary glands displayed similar low levels of *Rab6b* at L1 (Fig. 5C). These data showed that *Rab6b* remained poorly expressed at the onset of lactation and was not significantly affected upon *Rab6a* deletion.

### *Rab6a* loss in alveolar luminal cells leads to diminished STAT5 activation

The transcription factor, STAT5, plays a key role in secretory differentiation and activation of the alveolar epithelium, in particular downstream of prolactin receptor (PRLR) binding to PRL (Hennighausen and Robinson, 2008). To gain mechanistic insight into the defects observed in BlgCre; *Rab6a*^F/F^ mammary glands, we analyzed levels of the activated phosphorylated form of STAT5 (pSTAT5) in the protein extracts obtained at P18 and L1. WB data showed that compared to controls, mutant samples displayed a marked decrease in the ratios between pSTAT5 and total STAT5 amounts, indicating a reduced STAT5 activation upon *Rab6a* deletion (Fig. 6A). Expression of ELF5, a transcription factor controlling alveolar luminal cell maturation downstream of PRL signaling together with STAT5 (Lee and Ormandy 2012), was also diminished in mutant samples at L1 (Fig. S4D). On the other hand, similar levels of phosphorylated focal adhesion kinase (pFAK) were observed in mutant and controls samples at L1 (Fig. S4D). This indicated that integrin signaling, known to cooperate with PRL signaling for optimal STAT5 activation (Akhtar *et al.*, 2009), was not critically affected upon *Rab6a* deletion.

**Figure 6.**
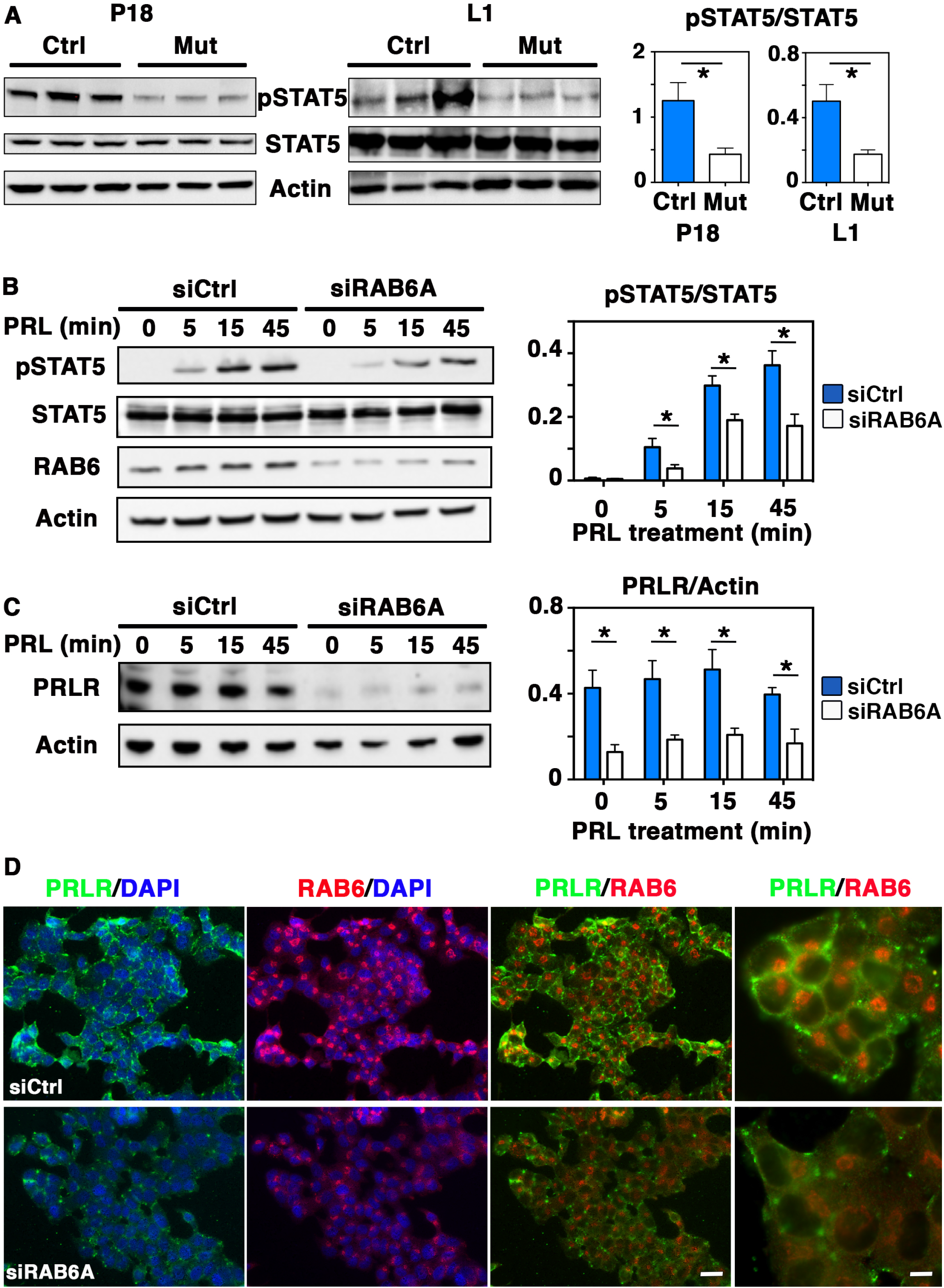
Decreased STAT5 activation in *Rab6a*-deficient mammary glands and RAB6A-depleted T47-D cells. **(A)** Left panels: western blots for pSTAT5, total STAT5 and actin performed on three distinct control and mutant mammary gland protein extracts at P18 and L1. Right panels: related quantification of pSTAT5/STAT5 amounts (mean ± SEM). *p=0.024 (P18), *p=0.045 (L1) **(B)** Left panel: Western blots for pSTAT5, total STAT5, RAB6 and Actin performed with T47-D cell lysates. SiCtrl and SiRAB6A cells were stimulated with PRL for 5, 15 and 45 min. Right panel: quantification of pSTAT5/STAT5 amount at each time point. Data are shown as mean ± SEM of 4 separate siRNA assays. *p<0.05 **(C)** Left panel: Western blots for PRLR and actin performed with the same T47-D cell lysates than in (B). Right panel: quantifications of PRLR amount normalized on actin at each time point. Data are shown as mean ± SEM of 3 separate siRNA assays. *p<0.05 **(D)** Immunolocalization of PRLR in SiCtrl and SiRAB6A T47-D cells. Enlarged areas of cells double-stained for PRLR and RAB6 are shown on the right. Bars, 30μm (left), 10μm (right).

To investigate the possible molecular mechanisms underlying the reduced STAT5 activation observed upon *Rab6a* deletion, we took advantage of T47-D, a human breast cancer-derived cell line commonly used for studying PRL-induced signaling events (Johnson *et al.*, 2010; Baker *et al.*, 2016; Oakes *et al.*, 2017). RAB6A depletion was achieved using siRNAs targeting *Rab6a/a’* variants, as reported (Patwardhan *et al.*, 2017). WB analysis of protein extracts showed that siRAB6A T47-D cells contained half of the RAB6 amount found in siCtrl T47-D cells (Fig. 6B and S4E). STAT5 being rapidly activated upon PRL binding, we analyzed T47-D cell response after 5, 15 and 45 min of PRL stimulation. Notably, at each time points, we observed a marked decrease in the pSTAT5/STAT5 ratios in RAB6A-depleted T47-D cells compared to non-depleted cells (Fig. 6B). To examine at which level PRL signaling was affected in this model, we analyzed PRLR expression using an antibody recognizing PRLR extracellular domain. Data from WB showed that RAB6A-depleted T47-D cells contained about half less PRLR than non-depleted cells and that the decreased PRLR amount persisted throughout PRL stimulation (Fig. 6C). Immunofluorescence studies indicated that RAB6A-depleted T47-D cells displayed less PRLR at their plasma membrane than non-depleted cells (Fig. 6D).

Overall, the results obtained with the mammary tissue and the T47-D cell line revealed that RAB6A depletion led to diminished STAT5 activation. In addition, they suggested that this defect probably occurred through an altered expression of PRLR at the cell surface, resulting in decreased PRL-induced signaling events.

## DISCUSSION

Our *in vivo* study uncovers a role for RAB6A in the differentiation and maturation of the luminal secretory lineage. Loss of *Rab6a* severely compromised the lactational function of the gland and the maintenance of the secretory tissue. It led to a decreased activation of STAT5, a key downstream effector of PRL-induced lactational processes. Data obtained with the PRL-responding human mammary cell line, T47-D, showed that loss of *Rab6a* led to decreased expression of PRLR, suggesting a perturbed transport of PRLR altering its surface expression and consequently its signaling cascade. These defects might largely account for the mammary phenotype observed in BlgCre; *Rab6a*^F/F^ females. Noticeably, despite the major functional deficiency of the RAB6A-depleted alveolar luminal cells, the polarized organization of the mammary bilayer was preserved *in vivo.*

### The luminal secretory lineage expresses a panel of *Rab* transcripts, including *Rab6a* and *Rab6a*’

Analysis of gene expression data revealed that the secretory lineage expresses a panel of *Rab* transcripts. These include the *Rab6a* and *Rab6a’* variants whose expression increased during pregnancy. In contrast, *Rab6b* remains poorly detectable in the secretory lineage in virgin as in the lactating gland. *Rab* genes encoding others “bona fide” Golgi-associated RAB GTPases, such as *Rab19*, *Rab30*, *Rab33b*, *Rab34*, *Rab36* and *Rab39b* (Goud et al., 2018), were absent from the secretory lineage, at least in virgin mice. *Rab* transcripts of GTPases involved in pre- and *cis*-Golgi (*Rab1a/b, Rab2a, Rab18)* as well as in late Golgi/TGN and post-Golgi (*Rab7*, *Rab8*, *Rab10*, *Rab11a*, *Rab14*) trafficking pathways were found significantly expressed in the secretory lineage.

### Loss of *Rab6a* in the secretory lineage has no major impact on mammary morphogenesis during gestation

To delete *Rab6a* in the secretory lineage we used the *Blg* promoter, known to be specifically active in the HR^-^ luminal progenitors and their secretory progeny from mid-pregnancy. This promoter is inactive in the basal and HR^+^ luminal cell lineages, throughout mammary development (Selbert *et al.*, 1998; Chapman *et al.*, 1999; Naylor *et al.*, 2005; Molyneux *et al.*, 2010; Di-Cicco *et al.*, 2015; Romagnoli *et al.*, 2020). The mammary epithelium expands massively till late pregnancy. During this morphogenetic period, the HR^-^ luminal and basal progenitors are amplified essentially through progesterone and local paracrine mediators secreted by the hormone-responsive HR^+^ cells (Fata *et al.*, 2000; Asselin-Labat *et al.*, 2010; Brisken and Ataca, 2015; Yu *et al.*, 2016). Whole mount and flow cytometry analyses did not reveal any severe underdevelopment of BlgCre; *Rab6a*^F/F^ mammary glands or perturbation of the luminal-to-basal cell balance, till late gestation. This indicated that basal and luminal progenitors have proliferated actively upon *Rab6a* deletion. Interestingly, unlike the control, the mutant epithelium was still actively proliferating at late gestation (P18), suggesting that loss of *Rab6a* may lead to a delayed mammary development possibly associated with an accumulation of HR^-^ luminal progenitors and perturbed differentiation events.

### RAB6A plays a crucial role in the differentiation, activation and maintenance of the luminal secretory cells

The major defects induced by loss of *Rab6a* clearly appeared at late gestation during the prelactogenic differentiation period and postpartum with the secretory activation. This led to a lactation deficiency in primiparous females, revealing a crucial role for RAB6A in the mammary gland function.

Mammary glands from numerous transgenic models showing deficiency in milk protein and lipid production display poor alveolar distension associated to defective alveolar luminal cell maturation (Naylor *et al.*, 2005; Li *et al.*, 2005; Russel *et al.*, 2011; Morales *et al.*, 2012; Akhtar *et al.*, 2016). This phenotype was observed in BlgCre; *Rab6a*^F/F^ at P18 and L1. Several aspects of the luminal secretory cell maturation were perturbed, in particular the global production of milk proteins, the expansion of CLDs and their apical targeting, and the baso-lateral expression of the glucose transporter, GLUT1. Conceivably, these perturbations altered both milk quantity and composition, compromising pup survival.

Loss of *Rab6a* led to a precocious luminal cell death as early as postpartum, showing that RAB6A is essential for the maintenance of the secretory tissue. The precise mechanisms remain to be determined however, it is known that the secretory differentiation and activation steps are critical to trigger and maintain the lactational process (Fata *et al.*, 2000; Vorbach *et al.*, 2002; Cui *et al.*, 2004; Anderson *et al.*, 2007; Russel *et al.*, 2011). When these steps are compromised, luminal cells enter into apoptosis, a physiological process that normally occurs after weaning and leads to mammary gland involution (Watson and Kreuzaler, 2011).

### Loss of *Rab6a* does not critically affect the polarized organization of the mammary bilayer

Loss of RAB6A during mouse embryogenesis and in migratory cell models has been reported to severely alter laminin and/or β1 integrin expression, perturbing cell adhesion and migration (Shafaq-Zadah *et al.*, 2016). In MDCK cells, a model presenting an apico-basal polarity in 3D cultures, RAB6A/B knock-out (RAB6-KO) strongly inhibited the secretion of soluble cargos, including laminin, whereas it mildly impaired that of transmembrane proteins (Homma *et al.*, 2019). Of note, despite their failure in laminin secretion, RAB6-KO MDCK cells were able to form polarized cysts in 3D cultures.

We did not detect any obvious perturbation of laminin deposition in the mammary epithelium from BlgCre; *Rab6a*^F/F^ females, at late gestation as during lactation. In the mammary bilayer, basal myoepithelial cells rather than luminal cells express a wide range of genes coding for extracellular matrix proteins (Kendrick *et al.*, 2008; Lim *et al.*, 2010). It is therefore considered that basal cells, together with the surrounding stromal cells, produce most of the basement membrane components of the mammary epithelium (Moumen *et al.*, 2011; Glukhova and Streuli, 2013).

Although less enriched in integrins than basal cells, luminal cells display several integrin chains, including α6 and β1 chains (Glukhova and Streuli, 2013; Romagnoli *et al.*, 2019 and 2020). Our data did not reveal notable perturbations neither in α6 and β1 chain expression nor in FAK activation in RAB6A-deficient luminal cells. Consistently, the mammary phenotype of BlgCre; *Rab6a*^F/F^ females did not phenocopy that observed in BlgCre; *Itgb1*^F/F^, BlgCre; *Itga3;Itga6*^F/F^ and BlgCre; *Ilk*^F/F^ mice (Naylor *et al.*, 2005; Akhtar and Streuli, 2013; Romagnoli *et al.*, 2020). In these models, during lactation, alveoli displayed luminal cell clusters protruding into the lumen, a phenotype absent upon *Rab6a* deletion.

Several essential markers of the apico-basal polarity, such as E-Cadherin, MUC1 and ZO-1, appeared properly addressed to the cell surface of RAB6A-deficient alveolar luminal cells. On the other hand, GM130 immunolabelling did not reveal mis-localization of the Golgi, a characteristic observed in BlgCre; *Itgb1*^F/F^, BlgCre; *Itga3;Itga6*^F/F^ and BlgCre; *Ilk*^F/F^ alveolar luminal cells that displayed perturbed apico-basal polarity (Akhtar and Streuli, 2013; Romagnoli *et al.*, 2020).

Potentially, other RAB proteins, highly expressed in the luminal secretory cells, could participate in maintaining apico-basal polarity features, in particular RAB8, a RAB GTPase implicated in the same post-Golgi transport pathways than RAB6A (Goud and Akhmanova, 2012). In addition, although expressed at very low level in the mammary tissue, RAB6B could have some compensatory function in the absence of RAB6A, as reported for RAB6-KO MDCK cells (Homma *et al.*, 2019).

### Loss of *Rab6a* led to a decreased activation of STAT5 in the secretory tissue

STAT5 is a key transcription factor controlling the mammary lobulo-alveolar development and the lactogenic function of the secretory lineage. Its activation level is strictly regulated in luminal cells: it gradually increases from mid-to late gestation and remains high throughout lactation (Hennighausen and Robinson, 2008; Hughes and Watson, 2012). Noticeably, we observed a decreased STAT5 activation in BlgCre; *Rab6a*^F/F^ mammary glands at P18 and L1.

Hennighausen and coll. have shown that STAT5 controls, directly or indirectly, the expression of multiple milk protein genes and regulatory molecules involved in the lactation process (Yamagi *et al.*, 2013). These include, in addition to the β-casein and *Wap* genes, *Adph* required for CLD maturation in the mammary tissue and *Rab18* implicated in CLD maturation and transport in lipogenic cells (Russel *et al.*, 2011; Dejgaard and Presley, 2019). STAT5 also controls lactose production by inducing the expression of α-lactalbumin and numerous members of the solute carrier family, such as *Slc2a1* encoding GLUT1 (Yamagi *et al.*, 2013). In addition, perturbation of STAT5 activation negatively impacts the expression of ELF5, a transcription factor cooperating with STAT5 in the specification of the secretory lineage (Lee and Ormandy, 2012; Yamagi *et al.*, 2013). Milk proteins, ELF5, ADPH, GLUT1 and RAB18 were found to display reduced or altered expression in BlgCre; *Rab6a*^F/F^ mammary glands at P18 and/or L1, indicating that deregulated STAT5 activation might largely account for the perturbed function of the RAB6A-deficient secretory tissue.

### Loss of *Rab6a* affects PRLR expression and its downstream STAT5 signaling

STAT5 is predominantly activated by PRL, a pituitary hormone that plays a major role during the pre-lactogenic and lactation periods (Hennighausen and Robinson, 2008; Hughes and Watson, 2012). Therefore, the decreased STAT5 activation observed at these stages in the mammary tissue from BlgCre; *Rab6a*^F/F^ females strongly evokes perturbation in the PRL/PRLR signaling events.

Studies on PRLR localization in the mouse mammary gland have been hampered by the lack of antibodies. To circumvent this difficulty, we used the well-characterized PRL-responding human T47-D cells (Johnson et al., 2010; Baker et al., 2016; Oakes et al., 2017). Using siRNAs, we found that RAB6A depletion in this model resulted in decreased PRLR amount and surface expression and in reduced STAT5 activation downstream of PRL action. RAB6A plays a role in all steps of post-Golgi secretion (Goud and Akhmanova, 2012; Fourrière *et al.*, 2019). Loss of RAB6A might thus perturb PRLR transport at different levels from Golgi to the plasma membrane, affecting its surface expression and thereby PRL-induced signaling events. Whether perturbation of intracellular PRLR transport leads to PRLR delivery to lysosomes for degradation, as reported for unsecreted cargos in RAB6-KO MDCK cells (Homma *et al.*, 2019), remains to be determined.

Our study reveals for the first time a role for RAB6A in the lactogenic function of the mammary luminal secretory lineage and suggests that RAB6A controls STAT5 activation by regulating PRLR surface expression. The level of PRLR expressed by mammary luminal cells is known to be critical for proper alveolar development and lactation, as observed in a mouse model containing a single *Prlr* functional allele and displaying haploinsufficiency (Ormandy *et al.*, 1997; Hennighausen and Robinson, 2008). In addition, high levels of STAT5 activation are required for luminal secretory cells to proceed through a full differentiation program (Yamagi *et al.*, 2013).

RAB6A has been implicated in matrix protein secretion, cell-matrix interactions and cell polarity in different models (Shafaq-Zadah *et al.*, 2016; Fourrière *et al.*, 2019; Homma *et al.*, 2019). In contrast, our data indicate that loss of *Rab6a* in the luminal secretory lineage had no major impact on the structure of the mammary tissue. RAB6A function therefore appears to depend on cell type specialization and tissue context. Conceivably, in polarized tissues, distinct polarity features-apico-basal, front-rear and dual-might influence RAB6A-dependent intracellular trafficking pathways.

## MATERIALS AND METHODS

### Mouse strains and transgenic mice

BlgCre transgenic mice, expressing the Cre recombinase under the control of the *Blg* promoter, were purchased from The Jackson Laboratory. *Rab6a*^F/F^ mice have been previously characterized (Bardin *et al.*, 2015). The care and use of animals were conducted in accordance with the European and National Regulation for the Protection of Vertebrate Animals used for Experimental and other Scientific Purposes (Facility license C750517/18).

### Dissociation of mouse mammary glands and flow cytometry analysis

Single cells were prepared from a pool of thoracic and inguinal mammary glands harvested from adult females (12-week-old virgin or 15- and 18-day-pregnant), as described in detail elsewhere (Di-Cicco *et al.*, 2015). Briefly, minced tissues were transferred to a digestion solution containing 3mg/mL collagenase (Roche), 100 units/mL hyaluronidase (Sigma-Aldrich) in CO2-independent medium (Gibco Life Technologies) completed with 5% fetal bovine serum (FBS, Lonza) and 2mM L-glutamine (Sigma-Aldrich), and incubated for 90 min at 37°C with shaking. Pellets of digested samples were centrifuged and successively treated at 37°C with solutions of 0.25% trypsin (Life Technologies) /0.1% versen (Biochrom) for 1 min, 5mg/ml dispase II (Roche)/ 0.1mg/mL DNAseI (Sigma-Aldrich) for 5 min. Pellets were treated with a cold ammonium chloride solution (Stem Cell Technologies) and filtered through a nylon mesh cell strainer with 40 μm pores (Fisher Scientific) before immunolabeling.

Freshly isolated mammary cells were incubated at 4°C for 20 min with the following antibodies from Biolegend: anti-CD24-BViolet421 (clone M1/69; #101826; 1/50), anti-CD49f-PeCy7 (clone GoH3; #313622; 1/50), anti-CD29-PeCy7 (clone HMβ1-1; #102222; 1/100), anti-CD45-APC (clone 30-F11; #103112; 1/100), anti-CD31-APC (clone MEC13.3; #102510; 1/100), anti-CD54-PE (clone YN1/1.7.4; #116107; 1/50). Labeled cells were analyzed and sorted out using a MoFlo Astrios cell sorter (Beckman Coulter). Data were analyzed using FlowJo software. Sorted cell population purity was at least 95%.

### Microarray data analysis of *Rab* expression

Global gene expression analysis was performed with total RNA extracted from luminal cell populations isolated by flow cytometry as previously reported (Chiche *et al.*, 2019). Samples were hybridized on Affymetrix GeneChip Mouse Genome 2.1ST arrays. Hierarchical clustering was performed using hclust (R) with Euclidean distance and Ward agglomeration method. GTPase genes were classified according to their expression level that was considered as significant if > 7.

### Whole-mount analyses and histology

Dissected mammary fat pads were spread onto glass slides, fixed in Methacarn (1/3/6 mixture of acetic acid/chloroform/methanol) overnight at room temperature and stained with carmine alum (Stem Cell Technologies), or fixed in 4% paraformaldehyde overnight at 4°C, as described elsewhere (Bresson *et al.*, 2018). For histological analyses, fixed glands were embedded in paraffin, and 7 μm-thick sections were cut, dewaxed and stain with hematoxylin-eosin. Image acquisition was performed using Nikon Eclipse 90i Upright microscope. ImageJ software (NIH) was used to quantify alveolar size.

Frozen sections (5 or 8 μm-thick) were obtained after embedding mammary glands into Tissue-Tek (Sakura), using a Leica Cryostat.

For RAB6 immunodetection, mammary glands were fixed in 4% paraformaldehyde overnight at 4°C, and after incubation time in sucrose solutions were embedded into Tissue-Tek (Sakura) before freezing.

### Immunohistofluorescence labeling

Paraffin sections were dewaxed, processed for acidic antigen retrieval, incubated overnight at 4°C with primary antibodies, and then at room temperature with secondary antibodies for 2 h.

Frozen sections fixed with 4% paraformaldehyde or acetone were incubated with antibodies as paraffin sections.

The following primary antibodies were used: anti-K5 and anti-K8 (Biolegend; #905501; 1/1000 and #904801; 1/100, respectively), anti-αSMA (Sigma Aldrich; #A2547; 1/200), anti-PR (Santa Cruz; #sc-7208; 1/200), anti-Ki67 (ThermoFisher Scientific; #MA5-14520; 1/100), anti-pan-laminin (Abcam; #ab11575; 1/100), anti-ZO-1 (Thermo Scientific; #61-7300; 1/200), anti-MUC1 (Abcam; #ab37435; 1/200), anti-E-cadherin ECCD-2 (Life Technology; #13-1900; 1/200), anti-adipophilin (Progen; #GP40; 1/100), anti-GLUT1 (Abcam; #40084; 1/50), anti-β1 integrin (Millipore; MAB1997; 1/100), anti-GM130 (BD Biosciences; #610823; 1/100), anti-RAB6 (Santa Cruz; #sc-310; 1/100). The anti-mouse β-casein was designed by Covalab.

Alexafluor-488 or Cy3 conjugated secondary antibodies were from Jackson ImmunoResearch Laboratories. Immunostained tissue sections were mounted in Prolong Gold antifade reagent with DAPI (Invitrogen, Life Technologies).

Image acquisition was performed using either a Leica DM 6000B microscope (Wetzlar, Germany) equiped with MetaMorph software, Nikon Confocal A1R microscope 60× CFI Plan oil objective (Apo VC/NA 1.4/WD 0.13), or an Inverted Laser Scanning Confocal LSM880NLO/Mai Tai Laser (Zeiss) with an Airyscan module (63x/1,4 OIL DICII PL APO (420782-9900)).

### TUNEL assay

For cell death analysis, glands were fixed in 4% paraformaldehyde in PBS, pH 7.5, O/N at 4°C. Dewaxed paraffin sections were analyzed for TdT digoxygenin nick-end labeling with Apoptag Plus (Sigma Aldrich) following manufacturer’s instructions. Methyl green was used as counterstaining. Image acquisition was performed using a Leica DM RBE optic microscope (Leica Microsystems GmbH, Allemagne) and number of apoptotic cells per field was count using ImageJ software.

### RNA extraction and RT-qPCR

RNA was isolated from whole mammary glands using Trizol reagent (Life Technologies) and further purified on a cleanup column (Qiagen). RNeasy Microkit was used for RNA extraction from isolated mammary cells, as described (Di-Cicco et al., 2015; Chiche et al., 2019). To avoid eventual DNA contamination, purified RNA was treated with DNAse (Qiagen). RNAs were reverse-transcribed using MMLV H(-) Point reverse transcriptase (Promega). Quantitative PCR was performed using the QuantiNova SYBR Green PCR Kit (Qiagen) on a LightCycler 480 real-time PCR system (Roche). The values obtained were normalized to *Gapdh* levels. The primers used for RT-qPCR analysis were purchased from SABiosciences/Qiagen or designed using Oligo 6.8 software (Molecular biology Insights) and synthesized by Sigma Aldrich. Primers for *Rab6a*, *Rab6a’* and *Rab6b* used in this study are listed in Table S1.

### T47-D cell culture and RAB6A knockdown by siRNA

The human mammary epithelial cell line, T47-D, was maintained at 5%CO2 in a 37°C incubator and grown in a phenol red free-RPMI 1640 medium containing 10% FBS and 5μg/mL insulin, as described (Baker et al., 2016).

For siRNAs knockdown experiments, cells were transfected the day after their seeding with siRNA controls (siLuciferase) or specific siRAB6A/A’ (50nM final concentration) in Lullaby reagent (OZ Biosciences). A second transfection was performed 24 hours after the first one. Experiments were conducted 72h after the last transfection.

24 hours before PRL treatment, cells were washed in PBS and medium was replaced by a serum-free medium. Cells were treated with human PRL (Sigma, #L4021) at 250ng/mL for 5, 15 and 45 min at 37°C in a medium supplemented with 5μg/mL insulin and 1μM dexamethasone, as described (Baker et al., 2016).

Immunofluorescence staining of PRLR (anti-PRLR, Invitrogen; #35-9200; 1/200). and RAB6 (AC306, produced and purified in B. Goud’s lab; 1/1000) were performed after fixing the cells with 4% paraformaldehyde and mild permeabilization with 0.2% saponin.

### Western blot analysis

Mammary tissue samples were homogenized in RIPA extraction buffer (0.1% SDS, 276mM NaCl, 40mM Tris pH7.5, 2% NP-40, 4mM EDTA, 20% glycerol, 1X proteases inhibitors) following further incubation during 20 min at 4°C in a rotation wheel. After centrifugation at 12000 rpm during 15 min at 4°C, supernatants containing extracted proteins were recovered and BCA Protein assay kit from Pierce (#23225) was used to estimate protein concentration. 40 μg of protein extracts where boiled for 5 min in Laemmli buffer before migration. Cell pellets were resuspended in 1x Laemmli buffer, vortexed and boiled for 5 min. Samples were run on NuPAGE Novex 4-12% Bis Tris gels (Life Technologies/Invitrogen) and transferred onto nitrocellulose. Membranes were incubated with 5% BSA in TBS containing 0.1% Tween 20 (TBST) for 1 hour at room temperature and with primary antibodies overnight at 4°C.

The following primary antibodies were used on mouse cell-derived lysates: anti-mouse milk proteins (Accurate Chemical; YNRMMSP; 1/2000), anti-mouse b-casein (a kind gift from C. Streuli, Manchester University), anti-RAB6 (AC306, produced and purified in B. Goud’s lab; 1/2000), anti-RAB5 (Cell Signaling Technology, CST; #3547S; 1/1000), anti-RAB8 (BD Biosciences; #610844; 1/1000), anti-RAB11 (BD Biosciences; #610656; 1/1000), anti RAB18 (Sigma; #SAB4200173; 1/1000), anti-GLUT1 (Abcam; #40084; 1/500), anti-ELF5 (Santa Cruz; sc-9645; 1/500), anti-STAT5 (Santa Cruz; #sc-1081; 1/10000), anti-pSTAT5-Tyr694 (CST; #9359; 1/1000), anti-pFAK-Tyr397 (CST; #3283S; 1/750), anti-FAK (CST; #3285; 1/1000). T47-D cell lysates were probed using anti-RAB6 (AC306, produced and purified in B. Goud’s lab; 1/2000), anti-STAT5 (BD Biosciences; #610191; 1/1000), anti-pSTAT5-Tyr694 (CST; #9359; 1/1000) and anti-PRLR (Invitrogen; #35-9200; 1/500).

Secondary antibodies coupled to horseradish peroxidase were from Jackson ImmunoResearch Laboratories. Detection was performed by chemiluminescence (Super signal West Pico+ or Femto, ThermoScientific). Quantitative analysis was performed with ImageLab.

### Statistical analysis

p-values were determined using Student *t*-test with two-tailed distribution and Welch’s correction, using the GraphPad prismv6 software. When specified, a Pearson’s Chi-square test was applied.

## ACKNOWLEDGMENTS

We are grateful to the personnel of the Animal facility (Céline Daviaud, Paul Bureau) and the Flow Cytometry Core facility (Annick Viguier, Sophie Grondin, Zosia Maciorowski, Coralie Guérin) at the Institut Curie. We greatly thank Amandine Di-Cicco (Institut Curie, CNRS-UMR144) for expert technical assistance; Pierre de la Grange (GenoSplice, France) for generating heatmap; Lucie Sengmanivong and Vincent Fraisier from the Cell and Tissue Imaging facility (PICT-IBiSA), Institut Curie, member of the French National Research Infrastructure France-BioImaging (ANR10-INBS-04). We also greatly thank Marina Glukhova and Vincent Goffin for valuable discussions.

