## Supplementary-Table and Figures for "RAB6 GTPase is a crucial regulator of the mammary secretory function controlling STAT5 activation"

**SUPPLEMENTARY MATERIALS**

**SUPPLEMENTARY TABLE**

| qPCR primers | Sequence 5'-3' |
|---|---|
| **Rab6a-F**<br>**Rab6a-R** | **GCC TCA TTC CCA GTT ACA TCC**<br>**TCC ATT TTG TAG TTT GCT GGA A** |
| **Rab6a'-F**<br>**Rab6a'-R** | **AAA CAA TGT ACT TGG AGG ATA GAA CC**<br>**CAA GCT CCT GAA CCG CTC T** |
| **Rab6b-F**<br>**Rab6b-R** | **GGT TGC CTG GTA GGT GTT GT**<br>**GCT GCG AAA ATT CAA GTT GG** |
| **Gapdh-F**<br>**Gapdh-R** | **CCA ATG TGT CCG TCG TGG ATC**<br>**GTT GAA GTC GCA GGA GAC AAC** |

**Table S1: Primers for qPCR**

**SUPPLEMENTARY FIGURES LEGENDS**

**Figure S1. Mammary phenotype of Blg-Cre; *Rab6a*$^{F/F}$ females at day 15 of pregnancy (P15)**

**(A)** Flow cytometry analysis of mammary cells isolated from control (*Rab6a*$^{F/F}$) and mutant (Blg-Cre; *Rab6a*$^{F/F}$) females at P15. Upper dot-plots: double CD24/CD49f (α6 integrin, α6Itg) immunostaining. Lower dot-plots: double CD24/CD29 (β1 integrin, β1Itg) immunostaining. The gated basal (Ba) and luminal (Lu) cell populations are indicated.

**(B)** Percentage of CD24+ epithelial cells (left) and ratio between the luminal and basal cell populations (right) calculated from the flow cytometry data obtained at P15. Data are the mean ± SEM of 3 distinct control and mutant cell preparations.

**(C)** Deletion of *Rab6a* and *Rab6a'* in luminal cells isolated from P15 females, estimated by qPCR. Data are shown as mean ± SEM from 3 distinct mutant and control cell samples. qPCR values were normalized on *Gapdh* and control values were set to 1. * $p<0.001$

**(D)** Whole mount and immunohistological analyses of mammary glands from control and mutant P15 females. Upper panel: carmine staining. Bar, 2mm. Middle and lower panels: double K5/K8 and Ki67/DAPI immunofluorescence labeling. Bar, 30μm.

**(E)** Immunolocalization of adipophilin in control and mutant mammary tissues at P15. Adipophilin expression is restricted to luminal cells in alveolar buds. Bar, 60µm.

**(F)** Immunodetection of progesterone receptor (PR) in control and mutant mammary tissues at P15. Left: double PR/DAPI immunofluorescent staining. Bar, 45µm. Right: Percentages of PR+ luminal cells. Data are the mean ± SEM of counting performed on sections through 3 distinct control and mutant mammary glands. About 1500 DAPI-stained nuclei were counted on each section.

**Figure S2. Mammary phenotype of Blg-Cre; *Rab6a*$^{F/F}$ females at day 18 of pregnancy (P18)**

**(A)** Quantification of RAB6 depletion performed on the western blot shown in Fig. 2F. RAB6 expression was normalized to actin. Data are shown as mean ± SEM from four distinct control and mutant mammary gland extracts. *p=0.024

**(B)** Double immunolabeling for K5/K8 (upper panels; bar, 48µm) and E-cadherin/SMA (lower panels; ECad/SMA; bar, 24µm) in control and mutant mammary epithelium at P18.

**(C)** Immunolocalization of apical markers in control and mutant mammary tissues at P18. Double labeling of ZO-1/DAPI (upper panels; bar, 25µm) and MUC1/DAPI (lower panels; bar, 35µm).

**(D)** Immunolocalization of β1 integrin (β1Itg) in control and mutant mammary epithelium at P18. Nuclei are stained with DAPI. Enlarged views of the delineated areas are shown in the right panels. β1 integrin is strongly expressed by basal myoepithelial cells and is also present at the luminal cell-cell contacts in both mutant and control tissues. Bars, 45µm (left) and 20µm (right).

**(E)** Flow cytometry analysis of mammary cells isolated from mutant and control mice at P18. Upper panels: Cytograms showing double staining for CD24 and α6Itg (CD49f). The gated basal and luminal cell populations with their respective percentages are indicated. Lower panels: histograms of α6Itg (CD49f, left) and β1Itg (CD29, right) expression in the gated luminal cell population.

**Figure S3. Mammary phenotype of Blg-Cre; *Rab6a*$^{F/F}$ females at day 1 and 6 of lactation (L1 and L6)**

**(A)** Alveolar size distribution in mutant and control epithelium at L1, estimated in pixels. Data are from a pool of measurements performed on two distinct histological sections from 4 control and 4 mutant mammary glands. Pearson's Chi-square test: p<0.001

**(B)** Survival of pups nursed by control and mutant primiparous dams at L6. Litters from 5 control and 11 mutant females were analyzed. *p<0.0001

**(C)** Views of hematoxylin/eosin-stained histological sections through control and mutant mammary glands at L6. Bar, 650µm

**(D)** Weight of pups nursed by mutant and control dams after 4 and 6 days of lactation. At least 15 pups per control and mutant dams were weighted. Data are shown as mean ± SEM. *p<0.05

**Figure S4. Mammary phenotype of Blg-Cre; *Rab6a*$^{F/F}$ females at day 1 of lactation (L1) and RAB6A deletion in T47-D cells**

(A) Double immunofluorescence staining for E-cadherin/ZO-1 in control and mutant mammary epithelium at L1. Enlarged merge images of the delineated areas are shown on right panels. Bars, 48µm (left) and 27µm (right).

(B) Immunodetection of MUC1 in control and mutant mammary epithelium at L1. Nuclei are stained with DAPI. Bar, 48µm

(C) Immunodetection of β1 integrin (β1Itg) in control and mutant mammary epithelium at L1. Nuclei are stained with DAPI. Enlarged images of β1Itg expression in the delineated alveoli are shown on right panels. β1Itg is strongly expressed by basal myoepithelial cells and is also present at the luminal cell-cell contacts in both mutant and control tissues. Bars, 36µm (left) and 23µm (right).

(D) Western blots for ELF5, actin, pFAK and total FAK performed on four distinct control and mutant mammary gland protein extracts obtained at L1. The related quantifications of ELF5/Actin and pFAK/FAK, shown as mean ± SEM, are in the right panels: *p=0.05 (ELF5); ns, not significant

(E) RAB6 expression in siCtrl and siRAB6A T47-D cells calculated from western blot analyses. An example of western blot is shown in Fig. 6B. Data are the mean ± SEM from 3 distinct siRNA assays with all time points pooled. *p< 0.0001

Fig. S1

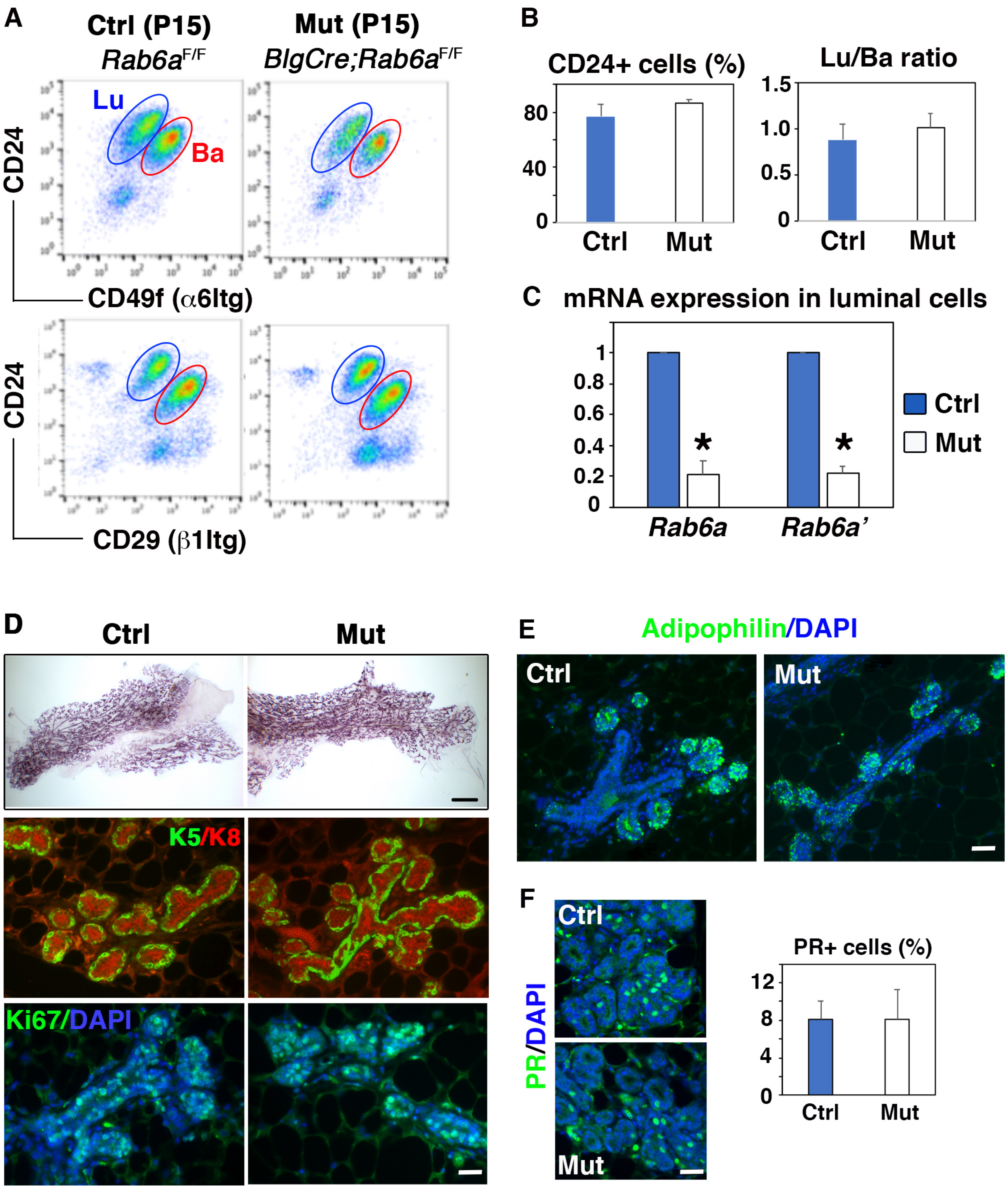
A
Ctrl (P15)
Rab6a F/F
Mut (P15)
BlgCre;Rab6a F/F
Lu
Ba
CD24
CD49f (α6Itg)
CD24
CD29 (β1Itg)
B
CD24+ cells (%)
80
40
0
Ctrl
Mut
Lu/Ba ratio
1.5
1.0
0.5
0
Ctrl
Mut
C
mRNA expression in luminal cells
1
0.8
0.6
0.4
0.2
0
*
*
Rab6a
Rab6a'
Ctrl
Mut
D
Ctrl
Mut
K5/K8
Ki67/DAPI
E
Adipophilin/DAPI
Ctrl
Mut
F
Ctrl
Mut
PR/DAPI
PR+ cells (%)
12
8
4
0
Ctrl
Mut

Fig. S2

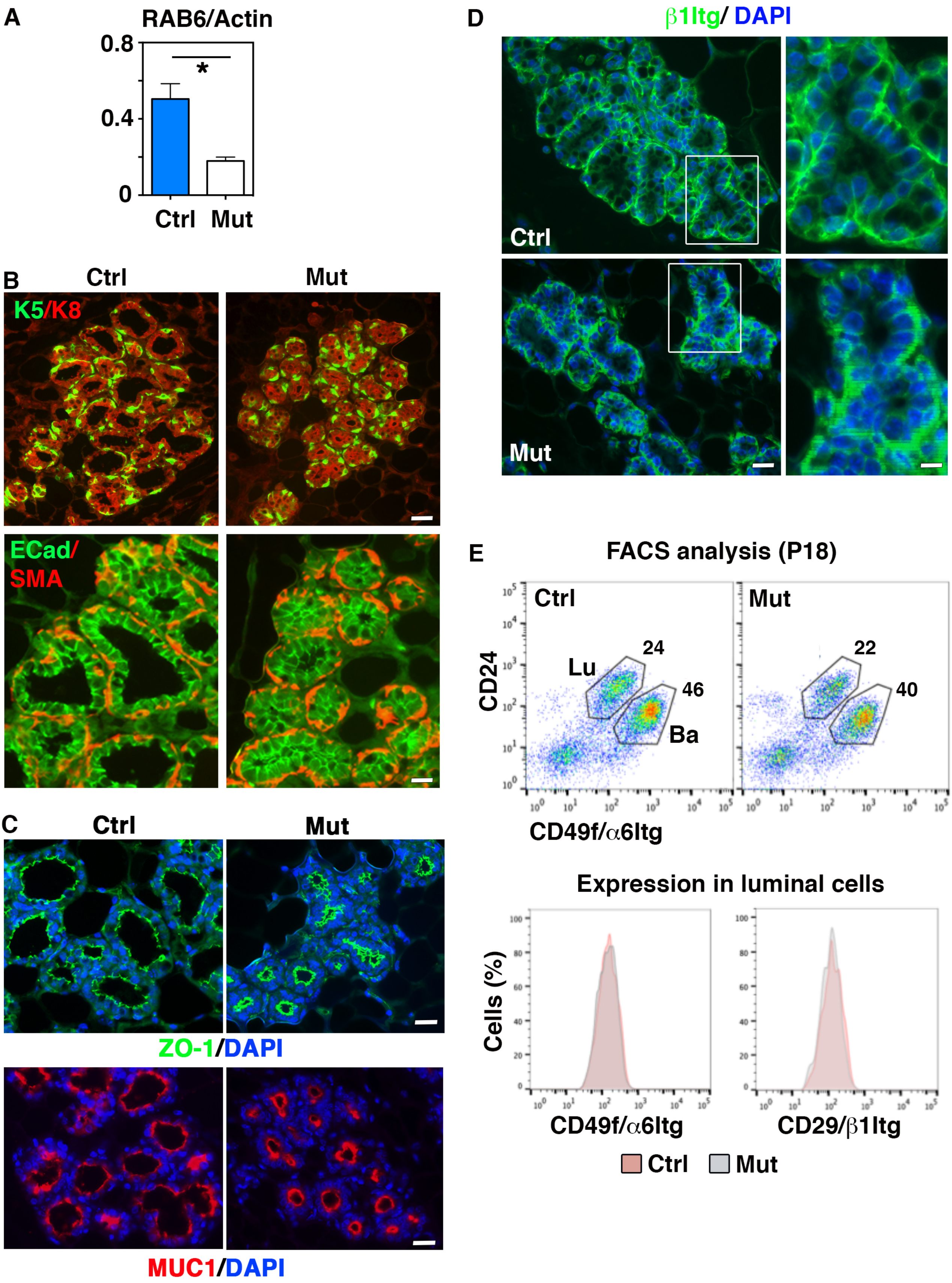
A
RAB6/Actin
0.8
0.4
0
*
Ctrl
Mut
B
Ctrl
Mut
K5/K8
ECad/
SMA
C
Ctrl
Mut
ZO-1/DAPI
MUC1/DAPI
D
β1Itg/ DAPI
Ctrl
Mut
E
FACS analysis (P18)
Ctrl
Mut
CD24
Lu
24
46
Ba
22
40
CD49f/α6Itg
Expression in luminal cells
Cells (%)
CD49f/α6Itg
CD29/β1Itg
Ctrl
Mut

Fig. S3

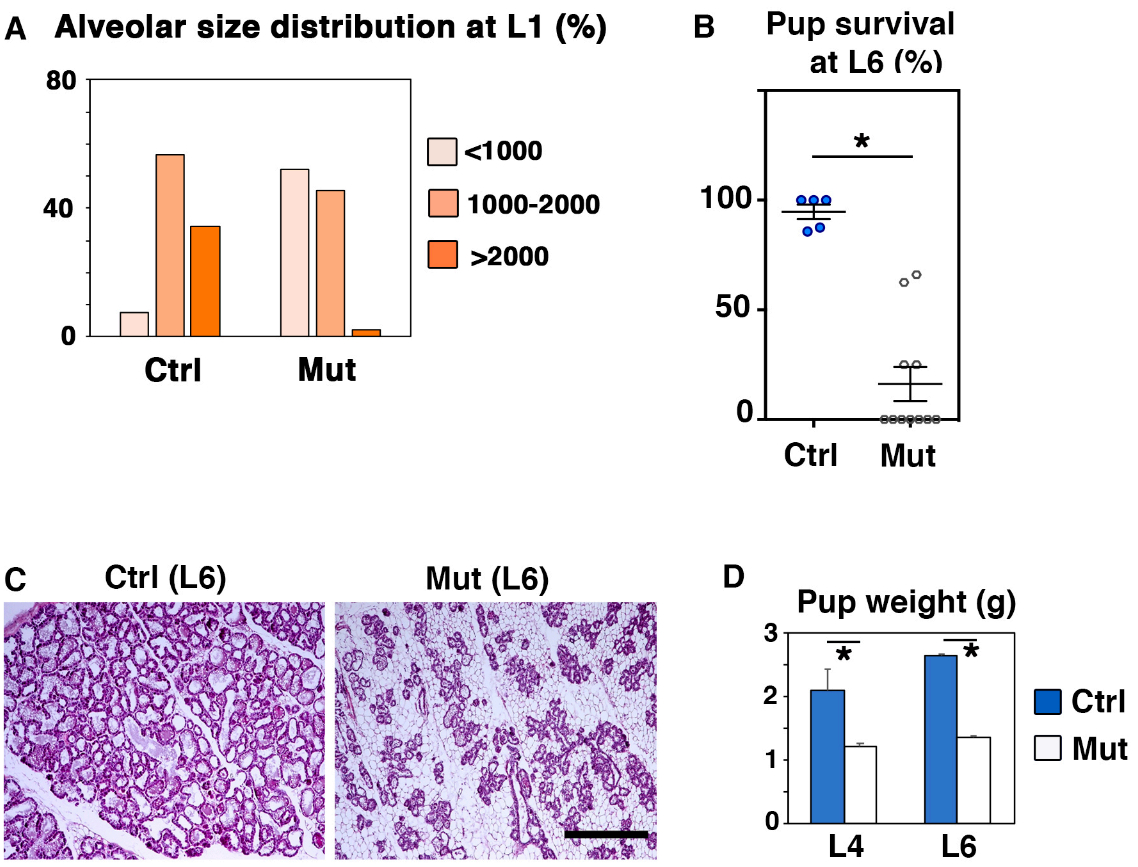
A
Alveolar size distribution at L1 (%)
80
40
0
Ctrl
Mut
<1000
1000-2000
>2000
B
Pup survival at L6 (%)
*
100
50
0
Ctrl
Mut
C
Ctrl (L6)
Mut (L6)
D
Pup weight (g)
*
*
3
2
1
0
L4
L6
Ctrl
Mut

Fig. S4

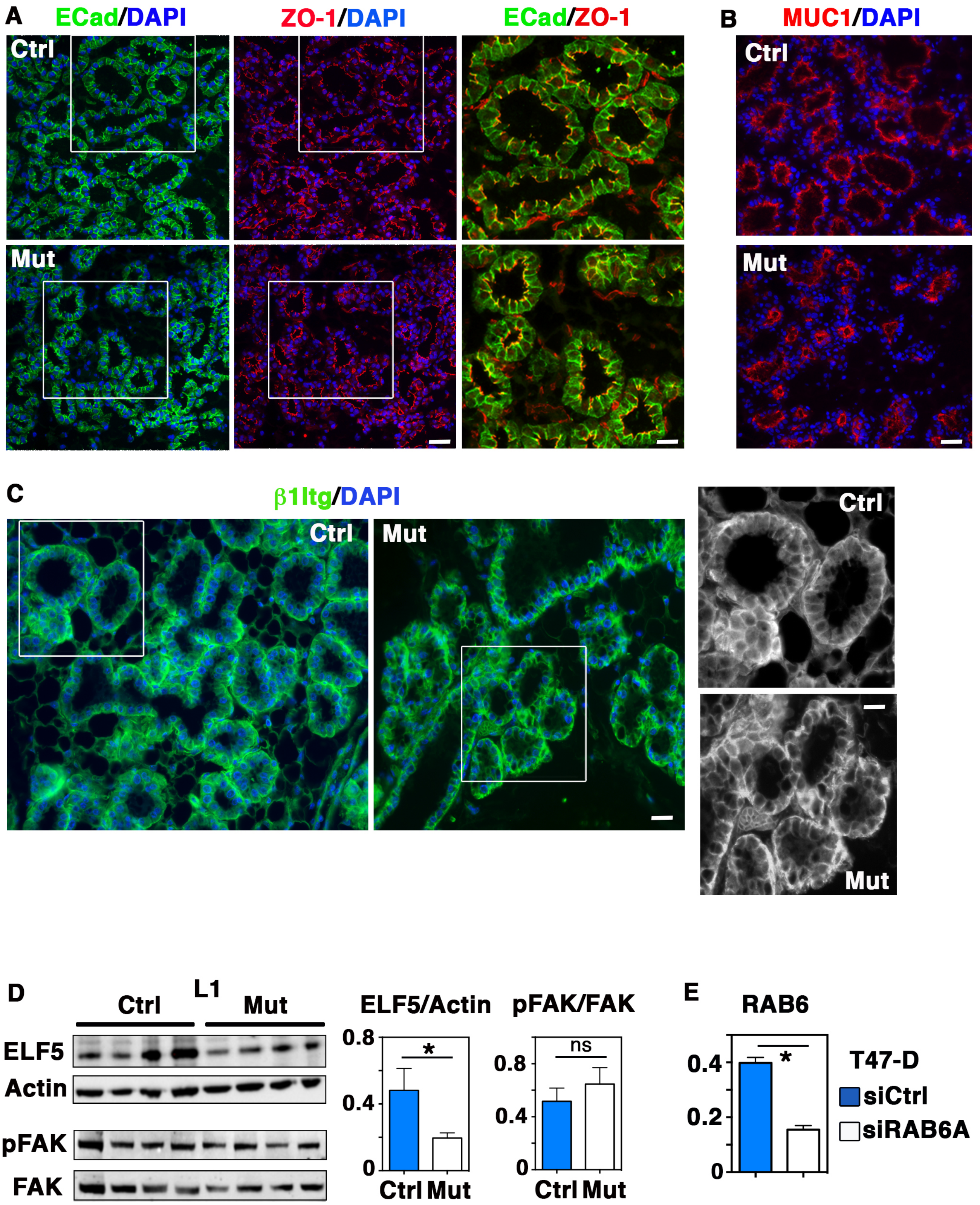
A
ECad/DAPI
ZO-1/DAPI
ECad/ZO-1
B
MUC1/DAPI
Ctrl
Mut
C
β1Itg/DAPI
Ctrl
Mut
D
Ctrl
L1
Mut
ELF5
Actin
pFAK
FAK
ELF5/Actin
0.8
0.4
0
*
Ctrl Mut
pFAK/FAK
ns
0.8
0.4
0
Ctrl Mut
E
RAB6
0.4
0.2
0
*
T47-D
siCtrl
siRAB6A
